# Antimicrobial activity of Mycobacteriophage D29 Lysin B during *Mycobacterium ulcerans* infection

**DOI:** 10.1101/507129

**Authors:** Alexandra G. Fraga, Gabriela Trigo, Juan Dominguez, Rita Silva-Gomes, Carine M. Gonçalves, Oliveira Hugo, António G. Castro, Joana Azeredo, Jorge Pedrosa

## Abstract

Buruli Ulcer (BU) is a cutaneous disease caused by *Mycobacterium ulcerans.* The pathogenesis of this disease is closely related to the secretion of the toxin mycolactone that induces extensive destruction of the skin and soft tissues. Although the World Health Organization recommends a combination of rifampicin and streptomycin for the treatment of BU, clinical management of advanced stages often requires extensive surgical resection of the infected tissue. Therefore, it is important to develop alternative strategies for the treatment of BU.

Endolysins (lysins) are phage encoded enzymes that degrade peptidoglycan of bacterial cell walls. Over the past years, lysins have been emerging as alternative antimicrobial agents against Gram-positive bacteria. Amongst Gram-positive bacteria, mycobacteria have an unusual outer membrane that is covalently attached to the mycolylarabinogalactan-peptidoglycan complex. To overcome this additional barrier to phage-mediated lysis, some mycobacteriophages encode a lipolytic enzyme, Lysin B (Lys B).

In this study, we demonstrated for the first time that recombinant Lys B displays lytic activity against *M. ulcerans* isolates. Moreover, using a mouse model of *M. ulcerans* footpad infection, we show that subcutaneous treatment with Lys B leads to a reduction in bacterial burdens, associated with IFN-γ and TNF production in the draining lymph node. These findings highlight the potential use of lysins as a novel therapeutic approach against this neglected tropical disease.

**AUTHOR SUMMARY:** Buruli Ulcer (BU) is a necrotizing skin disease caused by *Mycobacterium ulcerans.* Standard treatment for BU lesions consists of a combination of rifampicin and streptomycin for 8 weeks. However, clinical management of advanced stages of the disease often requires extensive surgical resection of the infected tissue. Therefore, it is important to develop alternative strategies for the treatment of BU. In that sense, we tested the efficacy of Lysin B (Lys B), a phage encoded lipolytic enzyme that degrades the mycolylarabinogalactan-peptidoglycan complex present in the mycobacterial cell wall. In this study, we show that Lys B not only displays lytic activity against *M. ulcerans* isolates *in vitro,* but also leads to a reduction in bacterial burdens in *M.* ulcerans-infected mouse footpads. These findings highlight the potential use of lysins as a novel therapeutic approach against this neglected tropical disease.

## INTRODUCTION

Buruli ulcer (BU) is a necrotic cutaneous disease caused by *Mycobacterium ulcerans* and represents the third most prevalent mycobacterial infection worldwide, after tuberculosis and leprosy [1-3].

BU’s pathogenesis is closely related to the secretion of the heat-stable polyketide toxin mycolactone that presents cytotoxic and immunosuppressive properties [4-8]. Early presentations of active BU include a painless pre-ulcerative nodule, papule, plaque or edematous lesion, which can evolve into typical ulcers or, in the most extreme cases, may result in extensive skin destruction, multifocal lesions or bone involvement [1,3,7,9,10].

The antibiotic regimen recommended by the World Health Organization (WHO) (daily administration of rifampicin (RIF) and streptomycin (STR) for 8 weeks) was proven to be curative for small pre-ulcerative lesions, but not for advanced forms, often requiring extensive surgical intervention [11-13]. Additionally, antibiotherapy is associated with adverse side effects, especially ototoxic and nephrotoxic effects [11,14].

Our group has previously demonstrated that bacteriophage therapy has potential as an innovative and effective therapy against *M. ulcerans* infection [15]. Indeed, our results in the murine model show that treatment with mycobacteriophage D29 effectively decreases the proliferation of *M. ulcerans* in the subcutaneous tissue resulting in marked macroscopic improvement of skin lesions [15].

Endolysins (lysins) are phage encoded enzymes produced during the late phase of the bacteriophage infection cycle, so as to degrade the cell wall peptidoglycan of the bacterial host, enabling the release of viral progeny [16-18]. Over the past decade, the development, characterization and exogenous application of recombinant and purified bacteriophage lytic enzymes has been successfully evaluated in several animal models of human diseases, such as sepsis, endocarditis, pharyngitis, pneumonia, meningitis and mucosal and skin infection [19-27]. Moreover, the use of a commercial endolysin for the treatment of *Staphylococcus aureus* skin infections has already been approved [28].

Amongst Gram-positive bacteria, mycobacteria have an unusual outer membrane composed by mycolic acids esterified with arabinogalactan (AG) which is linked to peptidoglycan, forming the mycolylarabinogalactan-peptidoglycan (mAGP) complex, a potential barrier to phage-mediated lysis [29]. Mycobacteriophage genome sequences show that, in addition to lysins that degrade the peptidoglycan layer of bacterial cell walls [30-32], some mycobacteriophages also encode a second lysin that targets the mAGP complex, known as Lysin B (Lys B) [33-35]. As described by Payne *et al.* [33], mycobacteriophage D29 encodes Lys B, a mycolylarabinogalactan esterase that cleaves the ester linkage joining the mycolic acid-rich outer membrane to AG, releasing free mycolic acids [36]. Although few studies have shown that Lys B can also act externally, suggesting its promising antimicrobial effect, there is no proven efficacy *in vivo* [37].

In the present study, following the *in vitro* evaluation of the lytic activity of recombinant mycobacteriophage D29 Lys B against *M. ulcerans* isolates, the therapeutic effect of Lys B during *M. ulcerans* infection was evaluated in the mouse footpad model of infection. The progression of macroscopic/microscopic pathology and bacterial loads, as well as the cytokine profiles, were evaluated in the footpad and the draining lymph node (DLN).

## MATERIALS AND METHODS

### Expression and purification of Lys B

The mycobacteriophage D29 Lys B gene (GenBank: accession number AF022214) was amplified by PCR from of mycobacteriophage D29 DNA using the primer: 5’- CCCTGGAACATATGAGCAAGCCC-3’ (nt 6606 to 7370). The amplification product was cloned into the expression vector pET28a (Novagem) fused to an N-terminal 6xHis tag. The resulting plasmid pET28a–Lys B was used to overexpress Lys B using *E. coli* BL21 (DE3) (Novagem) as a host for expression. Expression cultures were grown to an optical density (OD) between 0.4 and 0.6, at 600 nm, in Luria-Bertani broth containing kanamycin (50 μg/mL). Protein expression was induced with 1mM isopropyl-D-thiogalactopyranoside (IPTG) with shaking for 4 h at 37°C. Bacterial cells were harvested by centrifugation (10000x g, 5 min, 4°C), resuspended in phosphate buffer (50 mM NaH2PO4, 300 mM NaCl, 10 mM imidazole, pH 8), sonicated on ice for 5×10 s pulses separated by 10 s rests and then centrifuged (10000x g, 5 min, 4°C). The supernatant was applied to a nickel-nitrilotriacetic acid (Ni-NTA) agarose column (Qiagen) and the protein was eluted under native conditions with 500mM imidazole in phosphate buffer according to the manufacturer’s instructions. The purity of the protein was analyzed by 12% sodium dodecyl sulfate-polyacrylamide gel electrophoresis (SDS-PAGE) followed by Coomassie blue staining. Protein-containing fractions were combined and filter sterilized (0.22 μm pore size) for use in *in vitro* and *in vivo* studies. Protein concentration was determined using NanoDrop ND-1000 (NanoDrop Technologies). The lipolytic activity of the purified Lys B was confirmed by spotted in Middlebrook 7H9 plates containing substrates as follows: 1% (v/v) Tween 80 or 1% (v/v) Tween 20 with 1 mM CaCl2 and plates were incubated at 37°C for at least 24 h. Enzymatic activity was indicated by the formation of a white precipitate spot on Tween plates [34,38].

### *In vitro* Lys B activity against *M. ulcerans*

In order to evaluate the antimicrobial activity of purified Lys B against *M. ulcerans* strains, a plate lysis assay was performed [39]. Representative isolates of *M. ulcerans* from endemic BU areas were selected based on their genetic and phenotypic characteristics, including the type of mycolactone produced and their virulence for mice (see Table 1) [8,40-44]. The strains were obtained from the collection of the Institute of Tropical Medicine (ITM), Antwerp, Belgium.

**Table 1.**
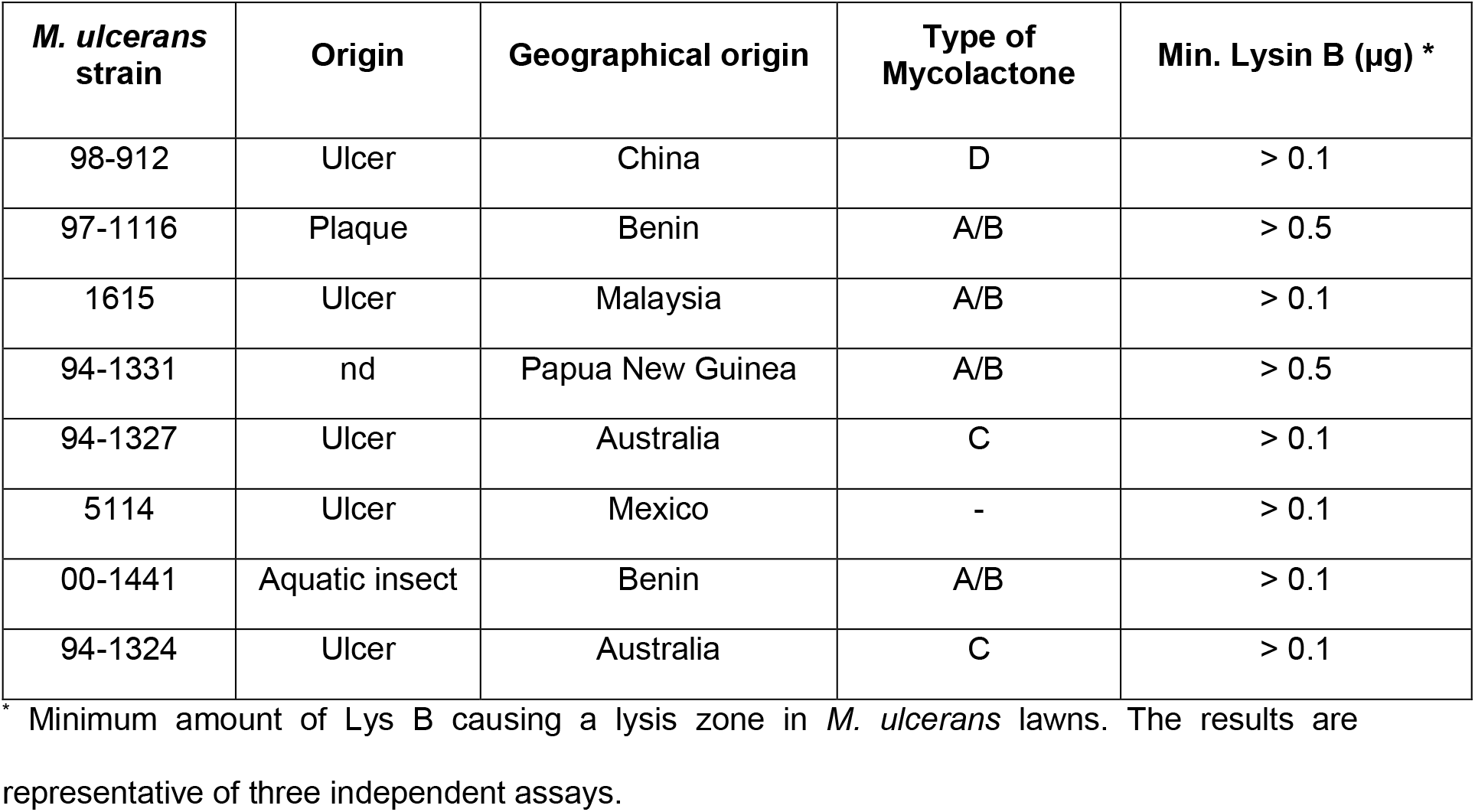
Antimicrobial activity of Lys B against *M. ulcerans* isolates.

Clinical isolates were grown on Middlebrook 7H9 broth medium at 32°C to an OD600 of 1.0 and clumps were dispersed by passing the bacterial suspension several times through a 25- gauge needle. The bacterial suspension (10^5^ CFU/mL) was plated on Middlebrook 7H9 agar medium. A stock solution of purified Lys B was serially diluted in phosphate buffer (final concentration 10-0.1 μg) and spotted onto bacterial lawns that air dried for 30 min. Phosphate buffer was spotted as a negative control. Plates were incubated at 32°C for approximately 6-8 weeks. Antimicrobial activity of Lys B was indicated by a clear lysis zone within the lawn where *M. ulcerans* growth was prevented.

### *In vivo* bioavailability and enzymatic activity of Lys B

The bioavailability and lipolytic enzymatic activity of Lys B were evaluated *in vivo.* Mice sera were collected by retro-orbital bleeding. Footpad and DLN suspensions were centrifuged for 10 min at 5000 rpm, and supernatant was considered the total protein extract. Protein concentration was determined using a NanoDrop ND-1000. The enzymatic activity of Lys B in samples was assessed by a lipase assay, as described above. For the bioavailability evaluation of the protein in samples, a western blot was performed. Briefly, samples were loaded and resolved by a 12% SDS-PAGE and transferred to 0.2 μm Nitrocellulose membranes (Bio-Rad) with the semi-dry Trans-Blot Turbo system (Bio-Rad). Membranes were blocked and subjected to immunoblotting with anti-mouse 6xHis antibody peroxidase conjugate (Clontech Lab. Inc., Takara). The 6xHis-tagged Lys B was detected with SuperSignal (Thermo Scientific #34095) in a Universal Hood II (Bio-Rad).

### Animals

Eight-week-old female BALB/c mice were obtained from Charles River (Barcelona, Spain) and were housed under biosafety level 3 conditions with food and water *ad libitum.*

### Footpad mouse model of *M. ulcerans* infection

*M. ulcerans* 98-912 is a mycolactone D producing strain, isolated in China from an ulcerative case and is highly virulent for mice, as previously described [8,42,43,45]. The isolate was grown on Middlebrook 7H9 agar medium at 32°C for approximately 6-8 weeks. For the preparation of inoculum, *M. ulcerans* was recovered, diluted in phosphate-buffered saline (PBS) and vortexed using glass beads. Mice were infected in the left hind footpad with 0.03 ml of *M. ulcerans* suspension containing 5.5 log_10_ CFU.

### Treatment of *M. ulcerans-infected* mice with Lys B

The treatment was initiated when footpads of mice were swollen to 2.7 mm and was performed by two subcutaneous (s.c.) injections in the infected footpad with 50 μg of Lys B in PBS at 10- and 13-days post-infection. Control-infected mice were injected with PBS without protein. Two groups of uninfected animals were also injected with Lys B or vehicle PBS buffer alone, as controls.

### Assessment of footpad swelling

After infection, as an index of lesion development, footpad swelling of mice was monitored throughout the experiment, using a caliper to measure the diameter of the frontal area of the footpad. For ethical reasons, mice were sacrificed after the emergence of ulceration and no further parameters were evaluated.

### Bacterial growth

*M. ulcerans* growth was evaluated in footpad tissues. Briefly, footpad tissue specimens were minced, resuspended in PBS (Sigma) and vortexed with glass beads to obtain homogenized suspensions. Serial dilutions of the footpad were plated on Middlebrook 7H9 agar medium. *M. ulcerans* numbers were counted after 6 to 8 weeks of incubation at 32°C and expressed as colony forming units (CFU).

### Detection of cytokines

The levels of the cytokines tumor necrosis factor (TNF) and gamma interferon (IFN-γ) in the supernatant of homogenized suspensions from DLN of mice were quantified by using a Quantikine Murine ELISA kit (eBioscience Inc), according to the manufacturer’s instructions.

### Detection of antibodies

The levels of antibodies against Lys B in the sera of mice were quantified by ELISA. Mice sera were collected by retro-orbital bleeding 3 days after the second s.c. Lys B injection. Briefly, polystyrene microtitre plates (Nunc) were coated with Lys B (10 μg/ml) and incubated overnight at 4°C. The wells were blocked with 0.05% (v/v) Tween 20 in PBS for 1h at room temperature. Serial dilutions of samples were plated and incubated for 2 h at room temperature. After washing, bound antibodies were detected with anti-mouse IgG antibody peroxidase conjugate (Sigma) for 1 h at room temperature. The antibodies against Lys B were detected with TMB substrate solution (eBioscience) at 450 nm. ELISA antibody titres were expressed as the reciprocal of the highest dilution giving an absorbance above that of the control (no Lys B injection).

### Ethics Statement

This study was approved by the Portuguese national authority for animal experimentation *Direção Geral de Alimentação e Veterinária* (DGAV 8421 from 2018). Animals were kept and handled in accordance with the Directive 2010/63/EU of the European Parliament and of the Council, on the protection of animals used for scientific purposes (transposed to Portuguese law - *Decreto-Lei* 2013/113, 7th of august).

### Statistical analysis

Differences between the means of experimental groups were analyzed with the two-tailed Student t test. Differences with a P value of ≤ 0.05 were considered significant.

## RESULTS

### Lys B shows *in vitro* antimicrobial activity against *M. ulcerans* isolates

Mycobacteriophage D29 Lys B was expressed and purified as a His6-tagged protein and its lipolytic activity was tested using Middlebrook 7H9 agar plates containing Tween 80 (Figure 1A) and Tween 20 (Figure 1B) as substrates [34]. Tween 80 and Tween 20 are esters of oleic (C18) and lauric (C12) acids, respectively, and can be cleaved by lipolytic enzymes to produce fatty acids and alcohol. The presence of Ca^2+^ in the medium causes the formation of an insoluble fatty acid salt that presents itself as a white precipitate [34,39]. As observed in Figure 1A-B, Lys B produced a spot of white precipitate, confirming the enzymatic lipolytic activity of purified recombinant mycobacteriophage D29 Lys B. No activity was observed with the phosphate buffer used as a negative control (data not shown).

**Figure 1.**
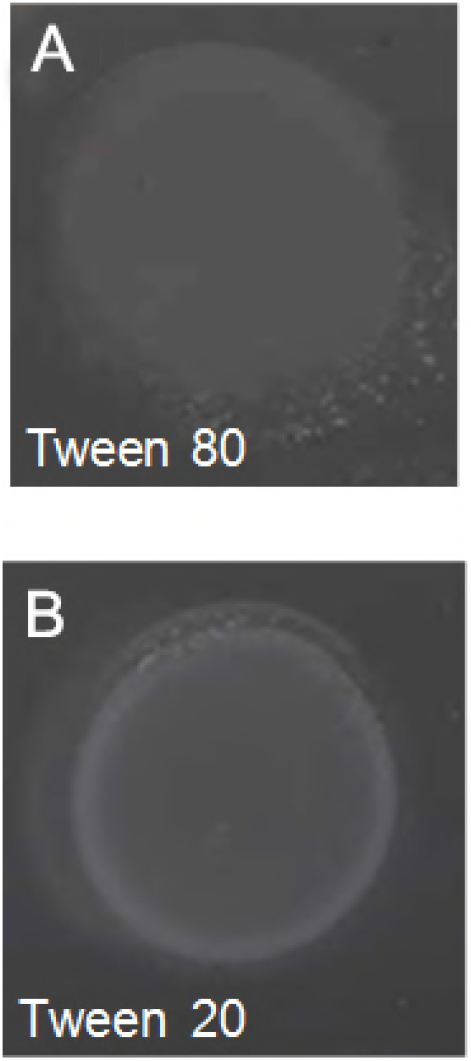
Lipolytic activity of Lys B. The lipolytic activity of the purified Lys B was confirmed by spotted in Middlebrook 7H9 plates containing substrates of (**A**) 1% (v/v) Tween 80 or (**B**) 1% (v/v) Tween 20 with 1 mM CaCl2. Plates were incubated at 37°C for at least 24 h. Enzymatic activity is indicated by the formation of a white precipitate spot on Tween plates

Purified Lys B was tested for its antimicrobial activity against several *M. ulcerans* isolates (Table 1) in a plate lysis assay. Our results show that Lys B has antimicrobial activity against *M. ulcerans.* Indeed, all *M. ulcerans* isolates tested were susceptible to the action of 10 to 0.1 μg of Lys B protein, causing a clear spot zone indicating cell lysis in *M. ulcerans* lawns. Moreover, only two of the tested *M. ulcerans* isolates, 97-1116 and 94-1331, were not susceptible to the lowest protein concentration used, 0.1 μg.

### Lys B is detected and has enzymatic activity *in vivo*

To determine the bioavailability and presence of enzymatically active Lys B *in vivo* after s.c injection in the footpad, the protein was measured in footpads, DLN, and sera. In a western blot analysis of footpad supernatant, Lys B was consistently detected, and the levels remained stable during the first 4h after administration (Figure 2A). At 6 h after injection, Lys B was still detected, although at lower levels (Figure 2A). In order to analyze if Lys B maintained its lipolytic enzymatic activity *in vivo,* samples were spotted on Tween plates [34,39]. As shown in Figure 2B, lipolytic activity was detected in footpad suspensions until 8h after injection. Regarding sera samples, although Lys B was not detected by western blot analysis (data not shown), lipolytic activity was observed (Figure 2B). No lipolytic activity or presence of Lys B was detected in the DLN (data not shown).

**Figure 2.**
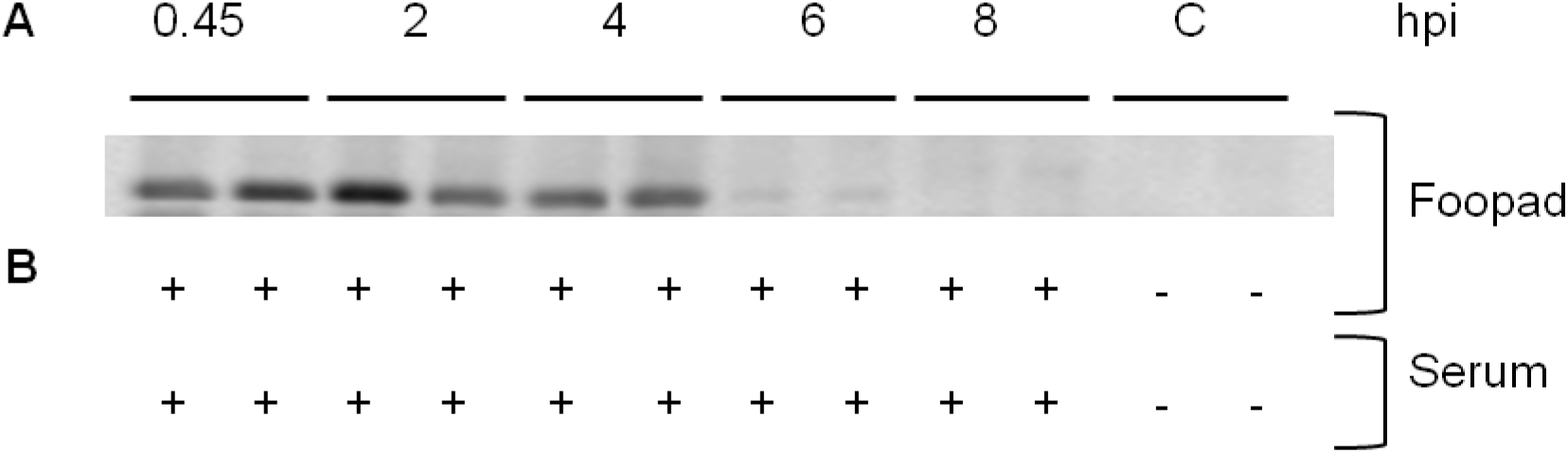
Assessment of bioavailability and enzymatic activity of Lys B in the footpad and serum of mice. Mice were injected subcutaneously in the left footpad with Lys B. At different time points, the presence of Lys B was assessed by Western blot (A) and Lys B enzymatic activity was determined by a lipase assay (B), in footpad supernatant and serum; hpi, hours post-injection; C, control mice; + lipase activity; - no activity. Results are from one representative experiment of two independent experiments.

### Treatment with Lys B decreases the *M. ulcerans* bacterial load in the footpad

To investigate the efficacy of Lys B treatment for the control of *M. ulcerans in vivo,* we used the footpad mouse model of infection [15,42-44]. Mice were s.c. infected in footpads with 5.5 log_10_ CFU of *M. ulcerans* strain 98-912. After 10 days, when footpad swelling reached 2.7 mm (Figure 3A), and at 13 days post-infection, mice were treated subcutaneously in the footpad with 50 μg of Lys B in PBS buffer. As shown in Figure 3B, *M. ulcerans* proliferated in infected footpads of non-treated mice over the course of experimental infection (P<0.01), while a significant 1log_10_ reduction in CFU counts (P<0.01) was observed in Lys B treated-mice at day 16 post-infection. The administration of vehicle PBS buffer or Lys B alone in uninfected footpads did not induce significant swelling of the footpad (data not shown). Moreover, our results show (Table 2) detectable levels of IgG antibodies to Lys B in sera of both Lys B-treated infected and non-infected mice at day 16 post-infection. Collectively, these results show that the administration of Lys B to *M. ulcerans* infected tissue has a protective effect, significantly reducing local bacterial burdens despite the presence of Lys B-specific antibodies in circulation.

**Figure 3.**
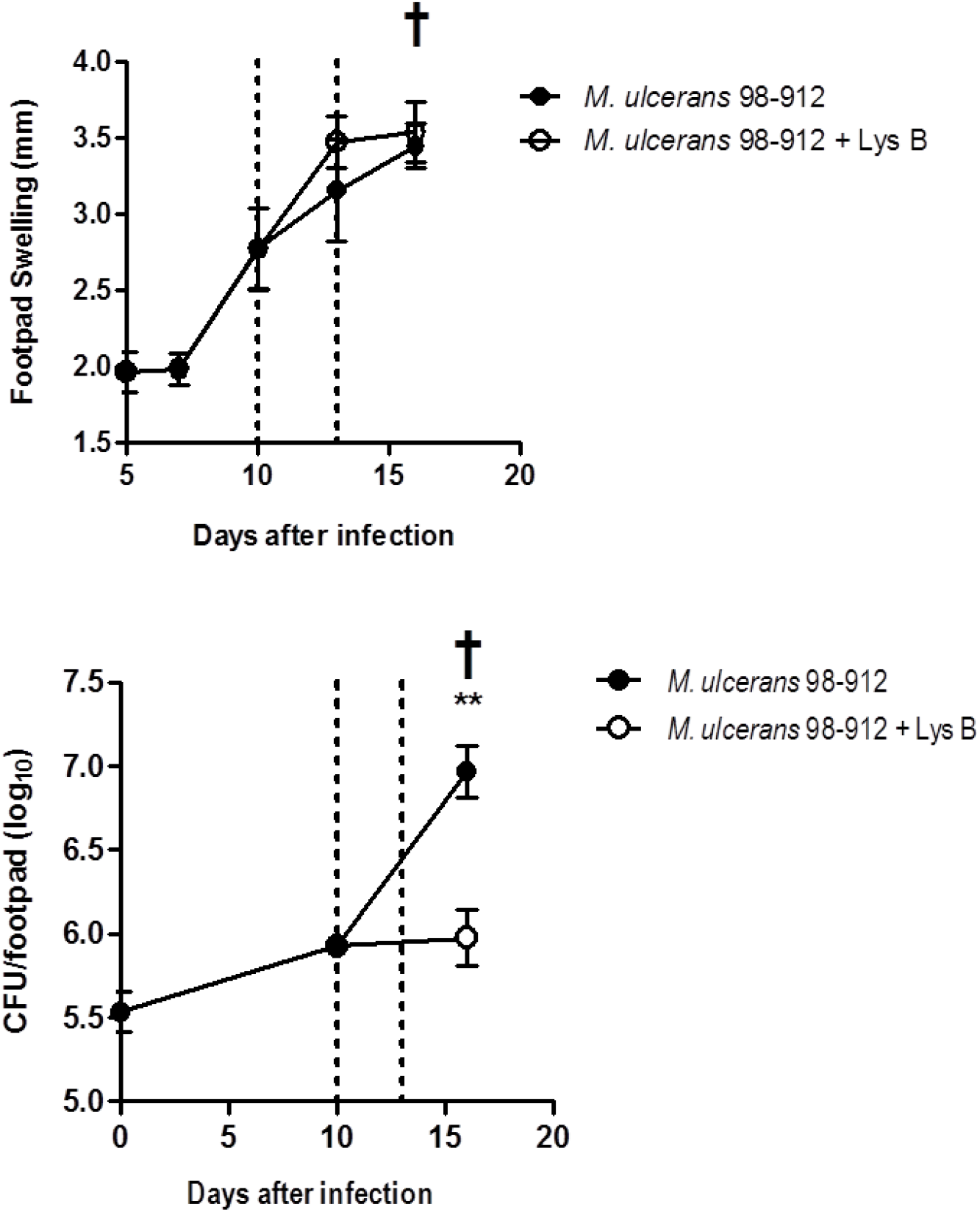
Lesion progression and *M. ulcerans* proliferation in footpads of infected mice. Mice were infected subcutaneously in the left footpad with 5.5 log_10_ CFU of *M. ulcerans* strain 98-912. After the emergence of macroscopic lesion (footpad swelling of 2.7mm) mice were subjected to treatment with two doses of subcutaneous injection of Lys B (10 and 13 days post-infection) (dashed lines). (**A**) Lesion progression was assessed by measurement of footpad swelling (n=15) and (**B**) bacterial proliferation was assessed by colony forming units in footpads (n=6). †, mice were sacrificed for ethical reasons after the emergence of ulceration. Results are from one representative experiment of two independent experiments. Data points represent the mean ± SD. Significant differences between treated and non-treated mice were performed using Student’s t test (**, p≤0.01).

**Table 2.** Antibodies titers against Lys B.

| Challenge <sup>a</sup> | Antibody titres <sup>b, c</sup> |
| --- | --- |
| *M. ulcerans* | ND |
| *M. ulcerans* + Lys B | 1166 ± 196 |
| Lys B | 1041 ± 165 |
| PBS | ND |

<sup>a</sup> Mice were injected subcutaneously in the left footpad with: *M. ulcerans* 98-912, Lys B or phosphate buffer. After the emergence of macroscopic lesion (10 days post-infection; footpad swelling of 2.7mm) infected mice were subjected to treatment with two doses of subcutaneous injection of Lys B (10 and 13 days post-infection).
<sup>b</sup> Results are from one representative experiment of two independent experiments; values represent the mean ± SD (n=6)
<sup>c</sup> ELISA IgG antibody titres are expressed as the reciprocal of the highest dilution giving an absorbance above that of the control (no Lys B injection).
ND, not detected

### Treatment with Lys B induces increased levels of IFN-γ and TNF in the DLN

Previous studies from our laboratory showed that *M. ulcerans* disseminates to the DLN, where the differentiation/expansion of mycobacteria-specific specific T cells occurs, contributing for the control of *M. ulcerans* proliferation through the production of IFN-γ [43,45] and TNF [8]. To determine whether Lys B treatment impacts host immune response, we carried out a comparative analysis of cytokine kinetics in the DLN.

Treatment with Lys B resulted in a significant increase in the levels of IFN-γ in the DLN (P<0.01) at day 16 post-infection (six days after the beginning of the treatment), as compared with non-treated mice (Figure 4A). The protein levels of the pro-inflammatory cytokine TNF were low in the DLN of non-treated mice (Figure 4B). In contrast, in Lys B-treated mice, significant levels of TNF (P<0.01) were detectable at day 16 post-infection (day 6 post-treatment) (Figure 4B).

**Figure 4.**
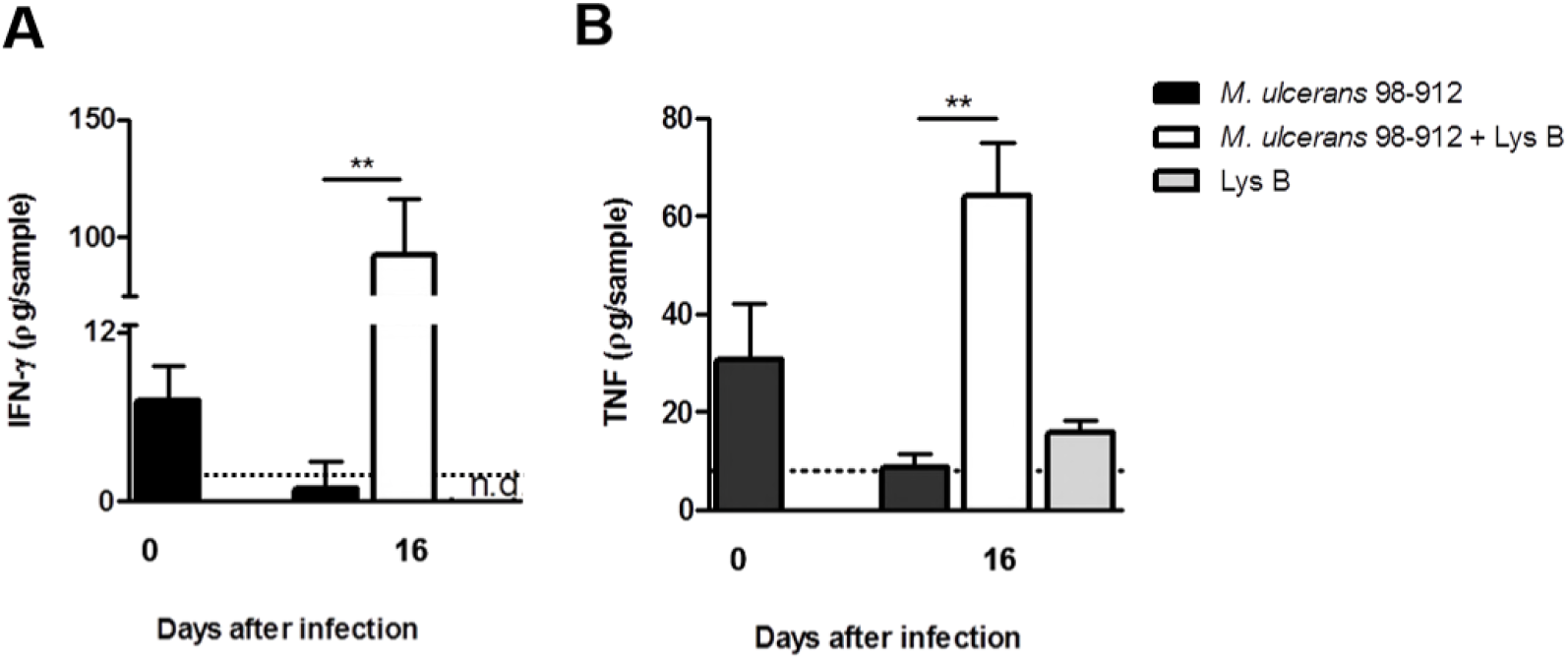
Cytokine profile in DLN of Lys B-treated *M. ulcerans* infected mice. Mice were infected subcutaneously in the left footpad with 5.5 log_10_ CFU of *M. ulcerans* strain 98-912. After the emergence of macroscopic lesion (footpad swelling of 2.7mm) mice were subjected to treatment with two doses of subcutaneous injection of Lys B (10 and 13 days post-infection). (**A**) Levels of IFN-γ and (**B**) TNF in DLN were quantified by ELISA. Results are from one representative experiment of two independent experiments. Bars represent the mean ± SD (n=6). n.d., not detected. Dashed lines represent the detection limit. Significant differences between treated and non-treated mice were performed using Student’s t test (**, p≤0.01).

## DISCUSSION

The RIF/STR regimen recommended by the WHO for the treatment of BU is often not effective in advanced ulcerative stages of the disease, for which the adjunction of surgical resection of infected skin followed by skin grafting is required [12,13]. Regarding the development of new and innovative anti-microbial strategies, we have previously shown in the mouse footpad model of *M. ulcerans* infection that a single s.c. injection of the lytic mycobacteriophage D29 can effectively decrease the proliferation of *M. ulcerans* resulting in marked macroscopic improvement of skin lesions [15]. Importantly, these results uncovered a second line of investigation. The potential of purified recombinant bacteriophage lytic enzymes (lysins) has been regarded as an alternative method to control bacterial pathogens, including *S. aureus* [23,24,39,46,47], *Streptococcus pneumoniae* [19,20,22,25,26], group B streptococci [24], *Bacillus anthracis* [48]. Recently, a new engineered endolysin (Artilysin Art-175) has shown activity against gram-negative pathogens, such as *Pseudomonas aeruginosa* [49]. In addition, endolysins have advantageous characteristics that avoid most of the common resistance mechanisms against antibiotics [48]. Although the ability of exogenously applied lysins as antibacterial agents is typically limited to bacteria without an outer membrane or surface lipids and waxes (i.e. Gram-positive bacteria), the mycobacteriophage Lys B has the potential to reduce cell viability due to the mechanical weakening and increased permeability of the cell wall. Lys B is a mycolylarabinogalactan esterase with activity in the mAGP, hydrolyzing the ester linkage that joins the mycolic acid-rich outer membrane to arabinogalactan, releasing free mycolic acids [33]. Due to the importance of the mAGP-complex for the stability of the mycobacterial cell envelope [50], we were prompted to test the therapeutic potential of Lys B in the context of *M. ulcerans* infection.

Herein, we have demonstrated the potential of mycobacteriophage D29 Lys B therapy against *M. ulcerans* infection in the mouse footpad model. Indeed, we show that treatment with Lys B can effectively decrease the proliferation of the high virulent mycolactone-producing *M. ulcerans* strain 98-912. Importantly, our *in vitro* results show a high susceptibility of several *M. ulcerans* isolates to Lys B, indicating that its activity *in vivo* is not limited to *M. ulcerans* 98-912. The therapeutic efficacy of Lys B treatment will depend on the presence of biologically active Lys B *in vivo.* As previously described, a rapid decrease in lysin levels after administration can result in the decrease of the amount of active lysin [22,51]. Indeed, in a pneumococcal meningitis mouse model, the bacterial load in the cerebrospinal fluid increased as the concentration of Cpl-1 lysin decreased over time [22]. Based on these observations, and in order to improve the Lys B bioavailability in mice footpads, we performed two administrations and shown detectable levels of Lys B protein with enzymatic activity in mouse footpads after s.c. injection.

It is known that the differentiation/proliferation of IFN-γ producing mycobacteria-specific lymphocytes occurs in mouse DLN early after experimental *M. ulcerans* infection [43]. However, this transient protective host response is not sufficient to inhibit the proliferation of virulent *M. ulcerans* in mice, as increasing concentrations of mycolactone at the infection site impair the effector activity of macrophages and induce cell and tissue destruction [8,42,45] In our study, Lys B administration effectively decreased the bacterial load in the infected tissue, allowing the development of a protective host immune response. Indeed, we observed that Lys B treatment in infected mice results in a significant increase in IFN-γ levels in the DLN.

Regarding antibody production, it has been demonstrated in several studies using different lysins and pathogens *(S. pneumoniae, S. aureus, Streptococcus pyogens* and *B. anthracis)* [52] that antibodies against lysins are non-neutralizing and therefore lysins can be used repeatedly to treat bacterial infections without adverse effects or loss of efficacy. In fact, when naïve and lysin-immunized mice were challenged with S. *pneumonia* and afterwards treated with Cpl-1 lysin, no differences were observed between the groups of mice regarding reduction of bacterial numbers [51]. Indeed, in the present study the presence of IgG antibodies against Lys B did not affect the control of viable bacteria in treated footpads, suggesting that produced antibodies did not inhibit Lys B activity.

For therapeutic use, an antimicrobial agent should not affect mammalian cells and only targeting the pathogen. Lysins target structures unique and highly conserved to bacterial cells and as such should not present a potential toxic threat to humans and animals [53]. This has been supported by several reports using lysins in preclinical studies in mouse models [26,51,54] and, in agreement, in our study, since Lys B injection was not associated with detectable side effects until the end of the experimental period.

To our knowledge, this is the first study on mycobacteriophage Lys B activity against *M. ulcerans* infection. It should be pointed out that mice were treated at an advanced stage of *M. ulcerans* infection, which is relevant for human infection since BU patients often seek medical treatment in advanced stages of the disease.

Although the development of a therapeutic regimen using Lys B will involve the determination of the proper doses and dosing schedules, our *in vivo* results are proof of concept of their potential against BU disease.

## Acknowledgments

We would like to thank the ICVS-Animal Facility for technical assistance with colony management and animal welfare. We would also like to acknowledge Leon Kluskens for technical assistance with cloning, expression and purification of Lys B.

